# Imputation of Low-density Marker Chip Data in Plant Breeding: Evaluation of Methods Based on Sugar Beet

**DOI:** 10.1101/2022.03.29.486246

**Authors:** Tobias Niehoff, Torsten Pook, Mahmood Gholami, Timothy Beissinger

## Abstract

Low-density genotyping followed by imputation reduces genotyping costs while still providing high-density marker information. An increased marker density has the potential to improve the outcome of all applications that are based on genomic data. This study investigates techniques for 1k to 20k genomic marker imputation for plant breeding programs with sugar beet as an example crop, where these are realistic marker numbers for modern breeding applications.

The generally accepted ‘gold standard’ for imputation, Beagle 5.1, was compared to the recently developed software AlphaPlantImpute2 which is designed specifically for plant breeding. For Beagle 5.1 and AlphaPlantImpute2, the imputation strategy as well as the imputation parameters were optimized in this study. We found that the imputation accuracy of Beagle could be tremendously improved (0.22 to 0.67) by tuning parameters, mainly by lowering the values for the parameter for the effective population size and increasing the number of iterations performed. Separating the phasing and imputation steps also improved accuracies when optimized parameters were used (0.67 to 0.82). We also found that the imputation accuracy of Beagle decreased when more low-density lines were included for imputation. AlphaPlantImpute2 produced very high accuracies without optimization (0.89) and was generally less responsive to optimization. Overall, AlphaPlantImpute2 performed relatively better for imputation while Beagle was better for phasing. Combining both tools yielded the highest accuracies.

**Summary:** Genotype marker information allows the prediction of an individual’s breeding value without the need to observe its actual phenotype which can accelerate the breeding progress. The more markers are genotyped, the better the genomic prediction may be. However, analyzing many markers is costly, particularly in commercial breeding programs where thousands of new individuals are genotyped. A solution to obtain information for all markers, while spending comparatively little on genotyping, is to genotype only a small fraction of markers in most individuals. Together with high-density information on other individuals, the low-density individuals can be imputed to high-density. High-density individuals are typically parents or highly influential individuals.

In this study, we compare the widely used software Beagle with the recently developed software AlphaPlantImpute2 on plant breeding data. To allow a fair comparison, we first optimized existing methods and developed new approaches. This was done to avoid comparing results of a less ideal version of one software to optimized settings of another software. After optimization, the software were evaluated in different scenarios with regards to genotyping errors, population types and number of markers based on simulated data. Simulated data were based on real marker data from a sugar beet population as input to mimic the population history of a commercial breeding population.

AlphaPlantImpute2 performs well with default parameters, while much optimization with regards to parameters and strategy was needed to boost accuracies of Beagle. A pipeline is presented which uses Beagle for phasing and AlphaPlantImpute2 for imputation. This pipeline yielded the highest accuracies and shortest run time.

**Core Ideas:** Beagle is sensitive to parameter tuning

Best imputation accuracies could be achieved by using a combination of Beagle and AlphaPlantImpute2

The population structure influence imputation accuracy

## 1 Introduction

Imputation allows the prediction of missing marker information in data sets. While it is not uncommon for a fraction of markers to have missing calls after genotyping, missing marker information can also be the result of merging two datasets that were genotyped with different marker panels.

Imputation is regularly used as a pre-processing step prior to genetic analyses. For applications that do not allow for missing values, including certain methods for genomic prediction (Meuwissen et al., 2001), imputing genetic raw data generated by SNP-chips or DNA sequencing is indispensable. It has also been shown to increase the detection power in Genome-Wide Association (GWA) studies (Nyine et al., 2019; D. R. Wang et al., 2018; Willer et al., 2008; Yan et al., 2017). Imputation also allows the aforementioned merging of data sets that are both genotyped with different marker panels assuming that a fraction of markers is genotyped in both sets. Geibel et al. (2021) showed how imputation of datasets comprised of sequenced and SNP chip genotyped animals can mitigate ascertainment bias which allows for more accurate estimation of population parameters.

Genotyping allows breeders to estimate genomic breeding values for new individuals at an early stage in a breeding program, thus reducing the generation interval. Basing selection on these ‘estimated breeding values’ can reduces the amount of resources and time otherwise spent on phenotyping (Windhausen et al., 2012). The accuracy of genomic prediction may increase with the number of individuals in the reference set and the density of markers genotyped albeit the improvement from genotyping additional markers diminishes the more markers are genotyped (Asoro et al., 2011; Erbe et al., 2013; Gorjanc, Battagin, et al., 2017; Hickey et al., 2014; Jacobson et al., 2015; Moghaddar et al., 2015; Norman et al., 2018). The rate of diminishing improvement depends on the population size, structure and diversity (Gorjanc, Battagin, et al., 2017; Lorenzana & Bernardo, 2009). For maintaining the genomic prediction accuracy across selection cycles, more markers are generally required to better capture the linkage disequilibrium structure (Asoro et al., 2011; Lorenzana & Bernardo, 2009; Norman et al., 2018).

A possible solution to obtain high-density marker data while spending comparatively little on genotyping is to deliberately genotype only a proportion of the population at high-density while genotyping the majority at low-density. The high-density individuals can then be used as a reference to impute the low-density individuals to high-density (Gorjanc, Battagin, et al., 2017; Gorjanc, Dumasy, et al., 2017; Jacobson et al., 2015; Thorn et al., 2021). Many imputation methods for plant and animal breeding have been developed over the years such as Fimpute (Sargolzaei et al., 2014), AlphaPlantImpute (Gonen et al., 2018), AlphaFamImpute (Whalen et al., 2020), AlphaImpute2 (Whalen & Hickey, 2020), OutcrossSeq (Chen et al., 2021), AlphaPlantImpute2 (Thorn et al., 2021), HBImpute (Pook et al., 2021). The majority of these methods are designed for genotyping only a fraction of the population at high-density. Often, these are the founders or parents that contribute all or most of the genetic material to the population. Generally, imputation infers missing genotypes based on reference haplotypes similar to the haplotype of the position in question. Offspring individuals can be explained as a mosaic of chromosome segments of a set of founders/ancestors. Since the breakage of haplotypes is relatively rare in breeding programs due to the low number of meiosis, parents make a powerful reference set.

Over the years, the software Beagle (Browning et al., 2018) has become a ‘gold standard’ for imputation; new software is typically tested against Beagle as a benchmark for performance. (Chen et al., 2021; Korkuć et al., 2019; Sargolzaei et al., 2014; Swarts et al., 2014; Thorn et al., 2021; X. Wang et al., 2020; Whalen et al., 2018; Whalen et al., 2020; Whalen & Hickey, 2020; Zheng et al., 2018). Beagle was originally developed for use in human genetics (Browning et al., 2018; Das et al., 2018) and its default settings are designed to work well for outbred human populations (Pook et al., 2020). Plant breeding populations, on the other hand, often show much higher levels of inbreeding and lower genetic diversity. Recently, Pook et al. (2020) demonstrated that Beagle imputation can be improved when working with data from breeding populations by tuning the imputation parameters. Thus, benchmarking the performance of Beagle in plant and animal breeding dataset requires adaptation of the parameter settings because the imputation performance in Beagle will otherwise be underestimated. This would lead to a biased comparison between approaches. Only a few studies use imputation parameters other than the default (Arouisse et al., 2020; Geibel et al., 2021; Lamb et al., 2021; Nyine et al., 2019; Thorn et al., 2021; Whalen & Hickey, 2020). These studies lower the parameter for the effective population size and some also increase the window size.

Thorn et al. (2021) recently presented a new method, AlphaPlantImpute2 (API2), and compared it to Beagle version 5.1. In this paper, we investigate the optimization of both tools further and give guidelines to researchers who want to improve their imputation accuracy. To allow for a fair comparison, we first optimized the imputation parameters and procedure for Beagle and AlphaPlantImpute2. We then tested the two tools on simulated populations based on real sugar beet marker data. While AlphaPlantImpute2 performed well with default settings, the accuracy of Beagle 5.1 improved four-fold when input parameters were optimized, the imputation approach was optimized, and pedigree information was used. We found that higher numbers of low-density offspring decreased accuracy when imputing with Beagle. As proof and as a solution to counteract this unwanted behavior, we suggest an approach which we call ‘single-genotype imputation’ which improves accuracies independent of the number of low-density lines. Furthermore, we observed that Beagle was superior at phasing, while AlphaPlantImpute2 was superior at imputation. We used the respective strength of both methods in a combined pipeline we call ‘Beagle + AlphaPlantImpute2’. We show its superiority in terms of run time and accuracy over other methods.

## 2 Materials and Methods

### 2.1 Raw data

To simulate the initial data to later compare imputation prediction to, we used proprietary real sugar beet marker data provided by KWS SAAT SE & Co. The data set contained bi-allelic marker information of 1655 lines, 41 of which are parents and the remainder are F3 offspring in 67 families. All lines were genotyped with a custom-made experimental SNP array that contains 19,519 markers covering all 9 chromosomes hereinafter referred to as ‘high-density (HD)’ or ‘20k’ panel. The parents showed various degrees of homozygosity. Together with a genetic map, three low-density panels containing 950, 1842 and 2719 markers were provided hereinafter referred to as ‘low-density (LD)’ or ‘1k/2k/3k’ panel, respectively. The smaller panels are subsets of the larger ones, i.e., all markers present on the 1k panel are also present on the 2k and 3k panel.

All lines with more than 10% missing marker information were removed from the original data set. Also, all lines with more than 100 Mendelian errors were removed. For this, the marker states of parents were assumed to be correct. After all filtering steps, 1357 lines from 36 families remained in the data set. 11 of the initial 41 parents were removed. About a quarter of all markers were monoallelic but not removed. The average fraction of missing marker information per genotype was 1.4% after all filtering steps.

### 2.2 Simulation of data sets

We phased and imputed the real 1357 lines with Beagle 5.1 (Browning et al., 2018) in one step with default settings except for the effective population size which was set to 30. After imputation and phasing by Beagle, no missing information was left in the data set. The phased parental information was then extracted and together with the provided genetic map and pedigree used for simulations with the R package ‘Meiosis’ (Müller & Broman, 2017). No mutations were induced. For traditional population types, the offspring were obtained by simulating the crosses as denoted in the pedigree followed by a number of selfings or double-haploid induction according to the population type simulated. Unless noted otherwise, five offspring per family were simulated. Bi-parental intercross populations were created by randomly selecting seeds from the F1 and allowing open-pollination within the family in the consecutive generations. Bi-parental intercross populations are denoted as e.g., OP10perfamn100 where 10 refers to the number of generations after the F1 while 100 refers to the population size. We also simulated open-pollinating populations based on all 30 founder lines. For open-pollinating populations, random mating started already from the parent generation on thus skipping the F1 creation step used for bi-parental populations. We denote these open-pollinating populations based on all 30 founders as e.g., OP10n100 with 10 referring to the number of generations and 100 to the family size in each generation. An OP10n100 is obtained one generation sooner than an OP10perfamn100 since no designated F1 generation is simulated. Table 1 provides an overview of the populations used in this study.

**Table 1:** Overview of populations used. The generations refer to the number of generations separating the population from the generation it is based on.

| Population | Based on | Mode of pollination | Generations | Population size | # families | Population type |
| --- | --- | --- | --- | --- | --- | --- |
| F1 | parents | cross | 1 | 5 per family | 36 | traditional |
| F2 | bi-parental F1 | selfing | 1 | 5 per family | 36 | traditional |
| F3 | bi-parental F1 | selfing | 2 | 5 per family | 36 | traditional |
| F10 | bi-parental F1 | selfing | 9 | 5 per family | 36 | traditional |
| DH | bi-parental F1 | / | 1 | 5 per family | 36 | traditional |
| OP10perfamn5 | bi-parental F1 | open-pollination within family | 10 | 5 | 36 | bi-parental intercross |
| OP10perfamn10 | bi-parental F1 | open-pollination within family | 10 | 10 | 36 | bi-parental intercross |
| OP10perfamn100 | bi-parental F1 | open-pollination within family | 10 | 100 | 36 | bi-parental intercross |
| OP100perfamn100 | bi-parental F1 | open-pollination within family | 100 | 100 | 36 | bi-parental intercross |
| OP10n5 | all 30 parents | open-pollination | 10 | 5 | 1 | open-pollinating |
| OP10n10 | all 30 parents | open-pollination | 10 | 10 | 1 | open-pollinating |
| OP10n100 | all 30 parents | open-pollination | 10 | 100 | 1 | open-pollinating |
| OP100n100 | all 30 parents | open-pollination | 100 | 100 | 1 | open-pollinating |

For comparability, we kept the number of lines constant for all population types, i.e., we increased or decreased the total number of offspring lines to 180 in the last generation of open-pollination.

Simulating datasets with different homozygosity levels for the parents was done by simulating open-pollination between all 30 parents thus also allowing selfing. One seed per parent was collected to replace the original parent mimicking female gametic control to reduce drift. These seeds were considered F1’s. Different parent population types like F2 or double haploids (DH) were created the same way as explained for the offspring. Whenever the original parents provided by KWS SAAT SE & Co were used in this study, they are denoted as ‘real’.

For simplicity, this study uses the term ‘line’ to refer to a genotypically distinct plant genotype regardless of its heterozygosity level. The parents of a line in this study are defined as those lines to who’s cross the line in question can be traced back. The line in question may be the result of the combination of the gamete of its parents (F1) or may be separated from the cross by any number of meiotic events like selfing.

### 2.3 Data set modification

Genotyping at low-density was mimicked by masking all markers not present on the low-density panel. The data of designated low-density lines was masked.

In a scenario investigating the influence of missing genotype calls, the markers at which to remove information were samples randomly per line. The fraction of missing markers was kept identical for all lines. In the scenario testing the influence of incorrect marker calls, we randomly sampled the markers per line at which the genotype state was to be changed. For every marker that was to be changed, we randomly set the genotype state to 0, 1 or 2 and never left the marker state as the true value. For example, if the true state is 0, only 1 or 2 were candidates. The fraction of errors induced prior to masking was identical for all lines. Every combination of factor levels was tested with five replicates. A new data set was simulated for every replicate. Unless noted otherwise, the data sets were simulated to mimic the following type:

- 30 real parents genotyped at high-density

- 36 families with 5 offspring per family

- F3 offspring genotyped with the 1k SNP panel

- No error induction

- No missing genotype call induction

### 2.4 Parameter optimization

To allow for a fair comparison between AlphaPlantImpute2 (Thorn et al., 2021) and Beagle 5.1 (beagle.18May20.d20), we first optimized the parameters for both tools. For AlphaPlantImpute2, this was done with a procedure in which only the high-density parent lines were used to create the haplotype library by setting ‘hd_threshold’ to 0.8. This library was then used for low-density offspring imputation without pedigree information. For parameter optimization in Beagle, we used a 2-step method in which all high-density and low-density data was imputed and phased by Beagle once. Then, only the phased information of the high-density lines was extracted and given as reference to a second Beagle run.

Parameters were optimized iteratively based on an incomplete grid search. The parameters were always set to be identical for both steps for all procedures using two steps. To derive parameters that are not different to default by chance, potentially better parameters were validated in all combinations with default parameters, except for the parameters ‘ne’ and ‘window’ of Beagle, as modifying these had been found to already have a large positive effect on accuracy.

### 2.5 Imputation procedures

After the optimized parameters were found, we tested different imputation procedures, or pipelines, that all utilized pedigree information.

To mimic the use of pedigree information in Beagle, a reduced reference panel that only contained the relevant parents was used, i.e., the parents of a family were looked up externally in R. Then, the reference set was restricted to only the parental lines of the family while the set that was to be imputed was restricted to only the offspring lines. This was repeated for all 36 families. We call this procedure ‘familywise imputation’. To check if imputation results can be improved further, we also did the phasing step familywise in addition to the familywise imputation step. To check whether running Beagle in two steps is really necessary, we also tried familywise imputation without the phasing step, i.e., no reference provided.

In AlphaPlantImpute2, we tested a procedure in which all high-density and low-density lines are used for the haplotype library creation step followed by imputation with pedigree information by specifying the founders file. We then tested a procedure in which only the parental lines were used for the haplotype library creation. In yet another procedure, to utilize pedigree information in the haplotype library creation step, we restricted lines specified for library creation to only lines of a family including both low- and high-density lines. Imputation was then also done per family and the process repeated for all families. This is comparable to the 2-step approach in Beagle where phasing and imputation are done familywise.

Based on our findings, we developed the single-genotype imputation pipeline for Beagle. Single-genotype imputation with Beagle is like the 2-step procedure without a pedigree as used for optimizing parameters, but instead of specifying all low-density lines in the second step, only one line is specified at a time. The procedure was repeated for all low-density lines. For the creation of the reference set, only high-density lines were considered. For single-genotype imputation, optimized parameters were used except for ‘iterations’, which was found to be sufficient when left at default.

The pipeline Beagle+API2 aims at combining the strengths of both tools. In this pipeline, only high-density lines are provided to Beagle in a first step. After they are phased, the lines are used as the haplotype library in AlphaPlantImpute2 for imputation of low-density lines. No pedigree information was utilized in either step. For Beagle+API2, the optimized parameters were used except for ‘iterations’ in Beagle which was found to be sufficient when left at default.

### 2.6 Scenarios

The following scenarios were tested with Beagle 5.1 and AlphaPlantImpute2:

1) Marker density on the low-density panel with the 1k, 2k and 3k panel

2) Various numbers (0, 1, 2, 3, 4) of the 5 offspring lines per family genotyped at high- density

3) Various fractions (0, 0.001, 0.005, 0.01, 0.02, 0.035, 0.05, 0.1) of missing genotype calls per line

4) Various fractions (0, 0.001, 0.005, 0.01, 0.02, 0.035, 0.05, 0.1) of genotyping errors per line

5) Different population types with every combination of offspring types with parent types offspring types:

F1, F2, F3, F10, DH,

OP10perfamn5, OP10perfamn10, OP10perfamn100, OP100perfamn100,

OP10n5, OP10n10, OP10n100, OP100n100

parent types:

real, F1, F2, F3, F10, DH

6) Imputing low-density parent genotypes based on 5 offspring per family with various numbers of offspring lines per family genotyped at high-density (1, 2, 3, 4, 5) and

7) Varying numbers of low-density offspring lines per family (2, 3, 4, 5, 7, 10, 15, 20, 25, 30, 35, 50)

For scenarios 1-5 and 7, imputation was done familywise with the 2-step procedure in Beagle. The procedure used with AlphaPlantImpute2 in these scenarios, used only high-density lines for haplotype library creation and the founder file was provided for imputation. In scenario 6, pedigree information was not used. Thus, the 2-step procedure without familywise imputation was used for Beagle. The founders file was not specified to AlphaPlantImpute2 for scenario 6.

The pipelines for single-genotype imputation with Beagle and Beagle+API2 were only tested on the following scenarios:

8) Imputing 5 low-density F3 offspring lines per family (36 families) based on all 30 real high-density parent lines

9) Imputing 50 low-density F3 offspring lines per family (36 families) based on all 30 real high-density parent lines

10) Imputing 5 low-density DH offspring lines per family (36 families) based on all 30 F1 high-density parent lines

11) Imputing 5 low-density F1 offspring lines per family (36 families) based on all 30 DH high-density parent lines

12) Imputing all 30 real low-density parent lines based 1 high-density F3 offspring line per family (36 families)

AlphaPlantImpute2 and Beagle were also tested on these scenarios (8-12). For AlphaPlantImpute2, the procedure considering only high-density lines in the haplotype library creation step was used. For Beagle, the 2-step procedure was used with the modification that the low-density lines were not used in the first step which is creating the reference set. In all scenarios, all pipelines were run without pedigree information.

### 2.7 Beagle versions

Two later versions of Beagle are available (5.2 and 5.3). We tried optimizing parameters for these two versions as well to not miss out on recent developments. The 2-step procedure as used in Beagle 5.1 before was used for version 5.2 and 5.3 too. Beagle version 5.2 (beagle.28Jun21.220), as presented by Browning et al. (2021), and Beagle version 5.3 (beagle.08Feb22.fa4) were used. For version 5.3, we always set the parameter ‘em’ to ‘false’.

### 2.8 Implementation

Custom R scripts were written for the imputation procedures, data handling and analysis. The imputation procedure to use, parameters and data set style were specified in custom made excel tables as input for the R scripts. R version 4.0.2 (R Core Team, 2020) was used. Computation was performed on the high-performance computing cluster of the Gesellschaft für wissenschaftliche Datenverarbeitung mbH Göttingen using Scientific Linux 7.9. The scripts and input tables for the imputation pipelines are available at https://github.com/tobiasniehoff/imputation_plant_breeding.

All imputation runs were allotted 8 GB of memory and 4 cores Broadwell Intel E5-2650 v4 with 2.2 GHz base frequency.

### 2.9 Performance measurement

The accuracy of imputation was measured as the Pearson correlation per line between the true simulated genotype states and the imputed genotype states. All markers on the high-density panel were considered. The reported figure in this paper is the average accuracy over all low-density lines. The run time was calculated as the difference of elapsed wall clock time between end and start of running the software, i.e., excluding all outside handling.

## 3 Results

### 3.1 Optimizing imputation

#### 3.1.1 AlphaPlantImpute2

Imputation accuracy achieved by AlphaPlantImpute2 with default parameters was 0.89 (leftmost procedure in Figure 1, plot B). This is one of the highest accuracies of all tested procedures in Beagle and API2, even when no pedigree information is supplied. The run time for default parameters with this procedure was about 360 minutes. Excluding low-density offspring from the reference library creation step drastically reduced the run time to 39 minutes while achieving an accuracy of 0.84 (procedure labeled as ‘parents for hap lib’ in plot B and D, Figure 1). We optimized parameters for this procedure. The optimized parameters used for AlphaPlantImpute2 imputation are presented in Table 2. With optimized parameters, this procedure yielded an accuracy and run time of 0.88 and 86 minutes, respectively. Supplying pedigree information and using high-density and low-density lines for library creation yielded accuracies of 0.92 and 0.91 and run times of 345 and 374 minutes for default and optimized parameters, respectively. Reducing the lines for the library creation step to only high-density lines lowered the accuracy for default parameters to 0.81 but optimized parameters maintained the accuracy at 0.9 (run times are 25 and 76 minutes, respectively) (procedure labeled as ‘parents for hap lib, imp w/ ped’ in plot B and D of Figure 1). This procedure with optimized parameters is used as the standard in this study (marked with ‘standard’ in plot B and D in Figure 1) When the library creation step and the imputation step are carried out per family, the accuracies for default and optimized parameters were both 0.81 while the run times were 210 and 260 minutes, respectively.

**Figure 1:**
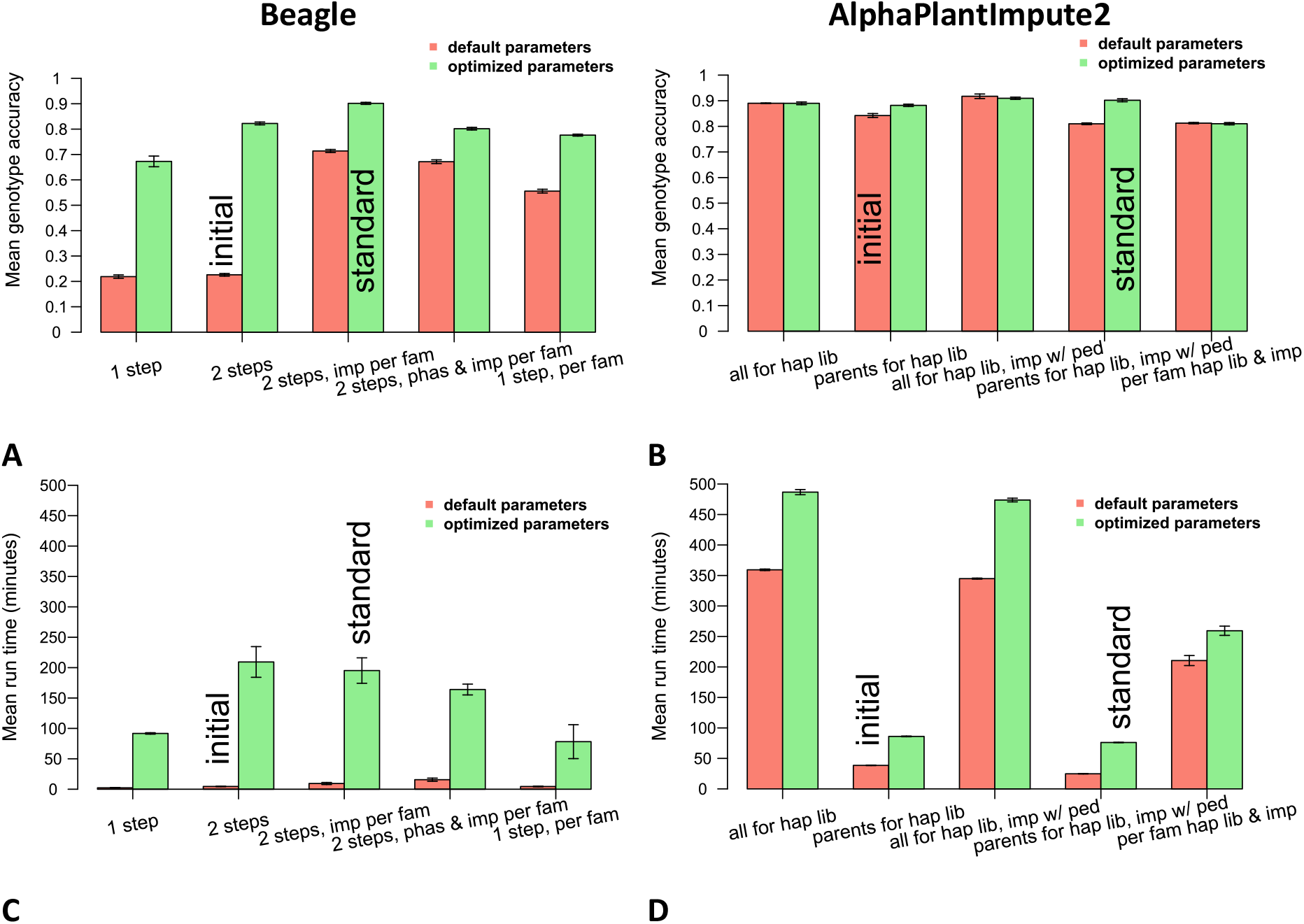
Accuracies and run times observed for imputation with default and optimized parameters as shown in Table 2 and Table 3 for different imputation procedures with AlphaPlantImpute2 and Beagle (each with 5 replicates). ‘initial’ marks the procedure used as a starting point for optimization. ‘standard’ marks the procedures used for all following tests unless mentioned otherwise. ‘2 steps’ refers to first running Beagle for phasing and then using the phased high-density parents as reference for imputation in a second step. ‘All’ and ‘parents’ for AlphaPlantImpute2 state what genotypes were considered for the creation of the haplotype library. ‘hap lib’ = haplotype library, ‘imp’ = imputation, ‘phas’ = phasing, ‘per fam’ = phasing or imputation restricted to only the genotypes and two parents of a family, ‘w/ ped’ = with pedigree/founders information.

**Table 2:** Parameters considered for finding optimized parameters for AlphaPlantImpute2. To only use high-density lines for the creation of the haplotype library, the parameter ‘hd_threshold’ was set to 0.8.

|  | default | range tested | optimized |
| --- | --- | --- | --- |
| **n_haplotypes** | 100 | 10-500 | 50 |
| **calling_threshold** | 0.1 | 0.01-0.7 | default |
| **n_sample_rounds** | 10 | 2-70 | 50 |
| **n_windows** | 5 | 2-20 | default |
| **error** | 0.01 | 0-0.1 | default |
| **recomb** | 1 | 0.1-2 | default |

**Table 3:** Values considered for finding optimal parameters in Beagle 5.1. Parameters not shown were not tested and their default values were used. The parameters were always identical for the phasing and the imputation step in the 2-step procedure. Imputation was not done familywise for optimizing parameters.

|  | **default** | **range tested** | **optimized** |
| --- | --- | --- | --- |
| **ne** | 1,000,000 | 4-1,000,000 | 4 |
| **window** | 40 | 1-400 | 100 |
| **burnin** | 6 | 1-1,000 | 90 |
| **iterations** | 12 | 2-20,000 | 5,000 |
| **phase-states** | 280 | 2-10,000 | 30 |
| **imp-states** | 1,600 | 2-5,000 | default |
| **imp-segment** | 6 | 0.1-700 | default |
| **imp-step** | 0.1 | 0.001-3 | default |
| **imp-nsteps** | 7 | 1-4,000 | default |
| **cluster** | 0.005 | 0.00001-10 | default |
| **overlap** | 4 | 0.01-20 | default |

### 3.1.2 Beagle

Compared to 1 step, running Beagle 5.1 in 2 steps to separate the phasing of parents from imputation of progeny did not change accuracies much when default paraments were used (0.22 for 1-step vs 0.23 for 2-step with run times of 2.3 and 4.5 minutes, respectively). With parameters optimized for the 2-step procedure, accuracies obtained with the 1-step procedure increased to 0.67 and 0.82 for the 2-step procedure with respective run times of 92 and 209 minutes. By doing the imputation step familywise in the 2-step procedure, i.e., restricting the set to be imputed to low-density sibling lines and restricting the reference to the two respective parents, the accuracy could be further increased to 0.9 for optimized parameters and 0.71 for default parameters with run times of 9.3 and 195 minutes, respectively. This procedure with optimized parameters is used as the standard in this study (marked with ‘standard’ in plot A and C in Figure 1). Doing the phasing and imputation step familywise yielded accuracies of 0.67 and 0.8 for default and optimized parameters, respectively. This procedure is considering the same information at every step as the AlphaPlantImpute2 procedure called ‘per fam hap lib & imp’.

Running Beagle in only one step familywise (rightmost pair of bars in plot A and plot C, Figure 1) resulted in accuracies of 0.55 and 0.78 for default and optimized parameters, respectively.

The optimized parameters for Beagle are presented in Table 3.

Note that although low-density genotypes were used for the creation of a phased reference set with Beagle, they were removed manually prior to supplying the reference set to Beagle in the second step, i.e., low-density lines were not provided as reference. The same was the case for AlphaPlantImpute2 whenever low-density genotypes are used in the haplotype library creation step.

We found that varying the parameter for the effective population size (‘ne’) had a strong effect on imputation accuracy with smaller values increasing the accuracy. No trend could be found for the population used by us for ne values in the range of 1 to 250 (plot A, Figure 2). However, the run time increased when decreasing values for the effective population size were used (plot B, Figure 2).

**Figure 2:**
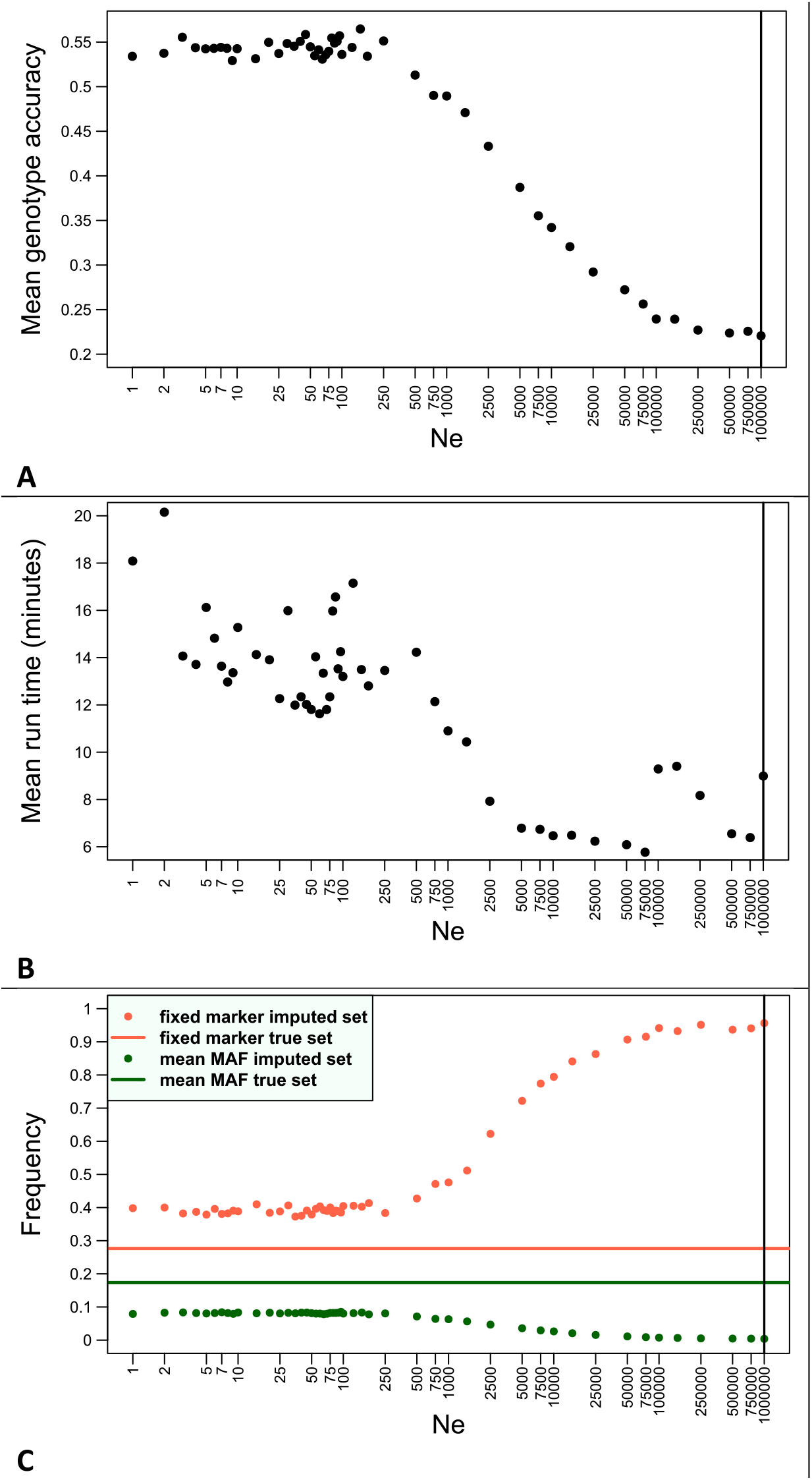
Effect of different values for ‘ne’on imputation accuracy (A), run time (B) average minor allele frequency and frequency of fixed loci (C) when imputing with Beagle. All other parameters are set as default. The imputation procedure used Beagle 5.1 in two steps without utilizing pedigree information (noted as ‘2 steps’ in plot A, Figure 1). Black line in A and B indicates default value (1,000,000). Plot C shows the mean minor allele frequency (MAF) and frequency of fixed markers in true and imputed data sets. Mean MAF and frequency of fixed (invariable) markers of true sets are shown as mean values over all sets for display purposes as the variance of these figures due to random mendelian sampling in F3 simulation is negligible.

Another Beagle parameter with a high impact on imputation accuracy is ‘iterations’. Increasing the number of iterations resulted in improved accuracies. For the data sets used in this study, no further increasing trend could be observed after about 3000 iterations (see plot A, Figure 3). The run time increased linearly the more iterations were used from about 8 minutes for 12 iterations (default) to about 690 minutes for 10,000 iterations (compare plot B, Figure 3).

**Figure 3:**
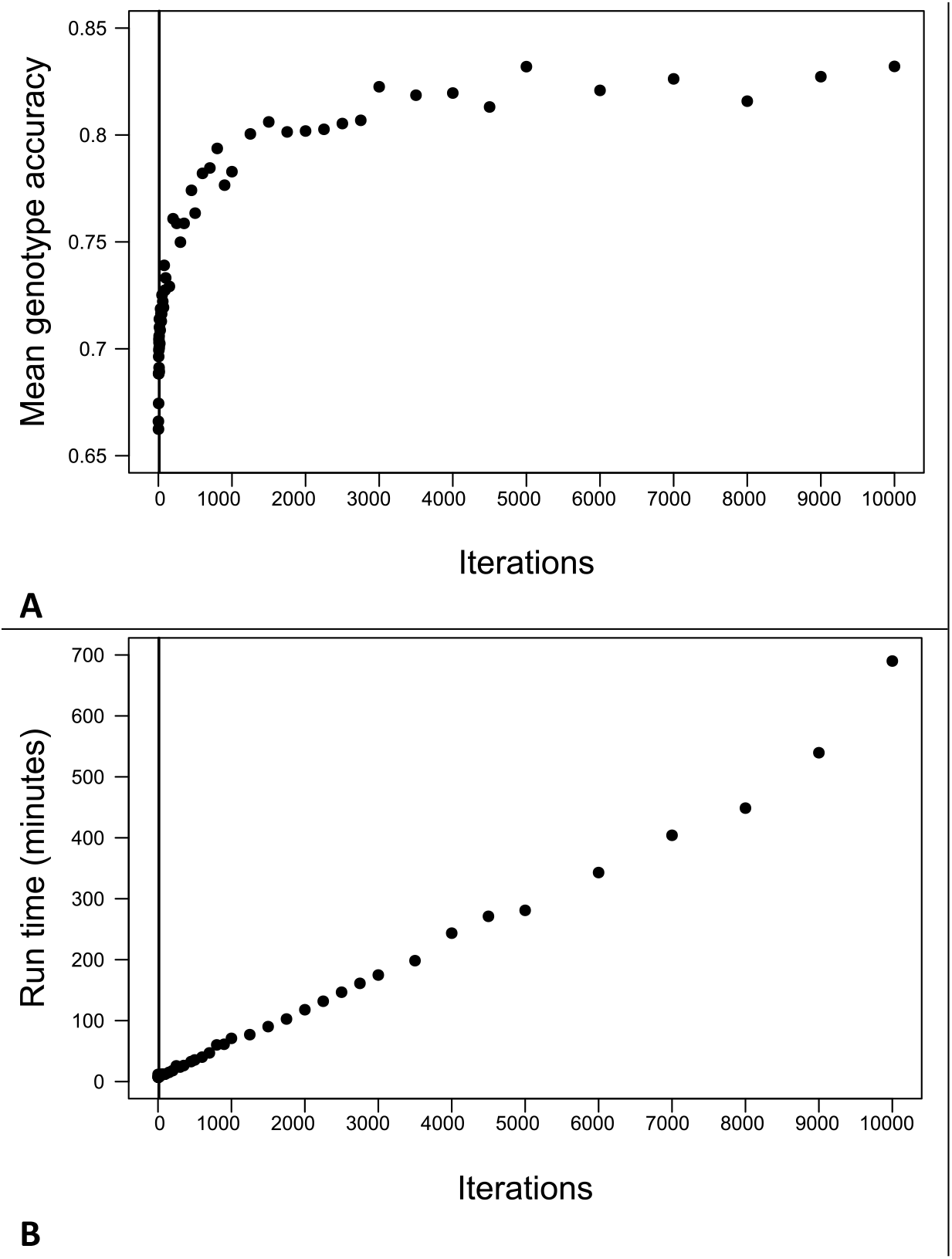
Effect of the number of iterations on imputation accuracy (A) and run time (B) of Beagle. For all other parameters other than ‘iterations’, the optimized values presented in Table 3 were used. The black line indicates the default value for iterations (12). The imputation procedure used Beagle in two steps without utilizing pedigree information (noted as ‘2 steps’ in plot A, Figure 1).

The MAF and the frequency of markers that are fixed were calculated in all tests conducted including the scenarios presented later in this study. Regardless of the software, the scenario and the imputation procedure or parameters used, the average MAF over all markers was always lower in the imputed set than in the simulated (true) set. The frequency of markers which are fixed, i.e., are invariable in the whole population (excluding high-density genotypes), was always found to be higher in the imputed set than in the true data set. With higher accuracies, both the mean MAF and the frequency of fixed markers converge towards the true value. These trends are visualized in plot C in Figure 2 for the example of decreasing ‘ne’ in Beagle.

### 3.2 Effect of marker density and genotyping offspring at high-density

Accuracies achieved with Beagle and AlphaPlantImpute2 could both be improved by including more markers on the low-density marker array (see Figure 4). Additionally, genotyping some offspring at high-density improved imputation accuracies. However, the additional gain in accuracy decreased the more offspring were already genotyped at high-density. AlphaPlantImpute2 was superior for the smallest low-density panel and the least high-density offspring whereas Beagle was relatively better for opposite cases.

**Figure 4:**
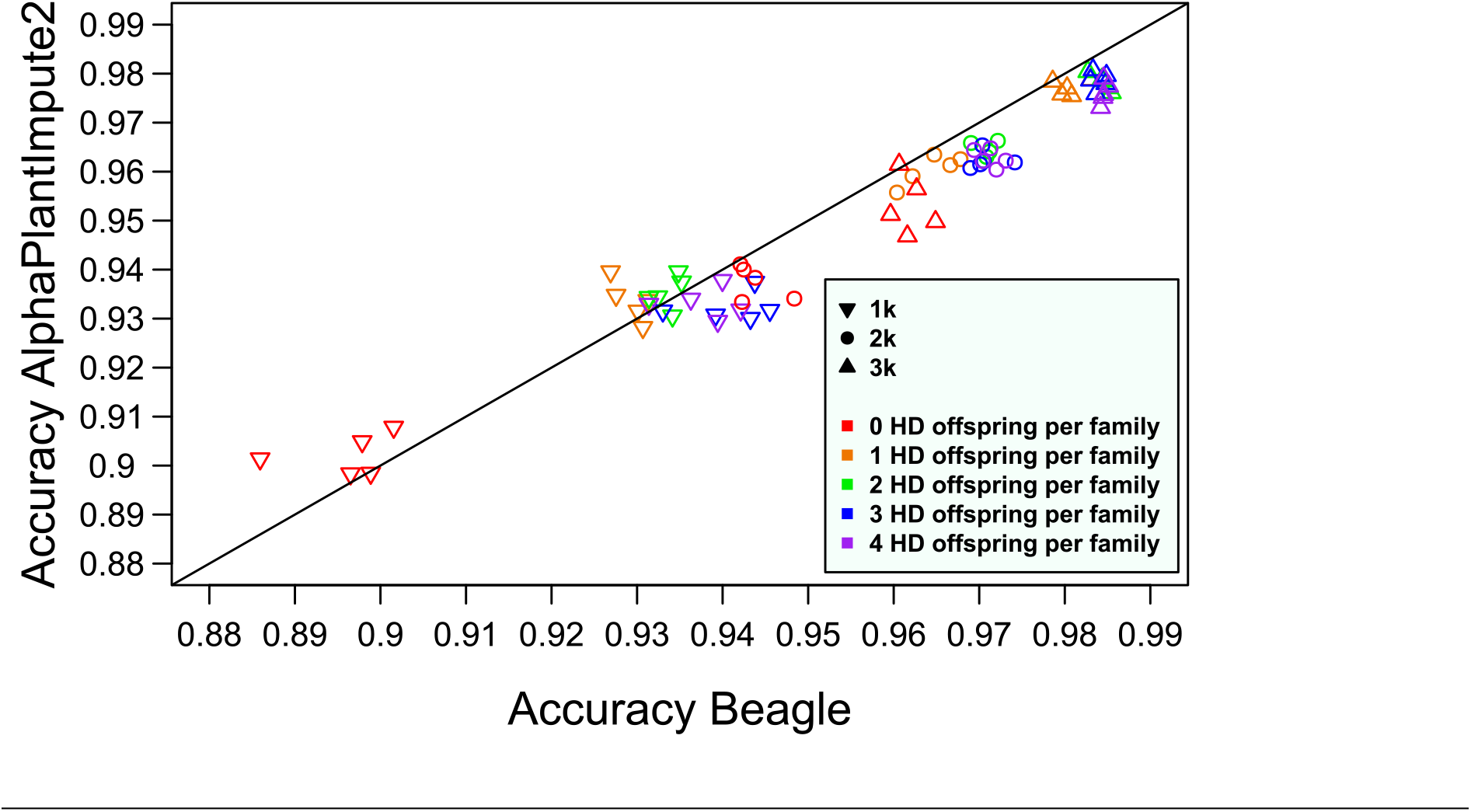
Effect of varying numbers of marker on the low-density panel and genotyping offspring at high-density (HD). Always five offspring per family were simulated of which some were genotyped at high-density. The accuracy is based on only low-density offspring.

Note that one high-density line per family means 36 high-density lines at the population level which can be used in the phasing step in the Beagle procedure. For AlphaPlantImpute2, the high-density offspring were included in the creation of the haplotype library in addition to the parents. For this scenario, the high-density offspring were included in the references of the families they were part of. Since the founders were specified for AlphaPlantImpute2, effectively only the two respective parents are used from the haplotype library for imputation whereas Beagle could sample haplotypes from all high-density family members.

The run time of AlphaPlantImpute2 increased by about 80 minutes for any increment increase in the number of high-density offspring. No trend in change of run time was observed for imputation with Beagle.

### 3.3 Missing genotype calls and genotyping errors

Both, AlphaPlantImpute2 and Beagle were able to make accurate imputation predictions when randomly placed missing genotype calls were induced in the data set for fractions of missingness lower than 2% (see Figure 5). At a missing rate of 0.1, the Beagle procedure achieved an average accuracy of 0.89 while AlphaPlantImpute2 yielded 0.86.

**Figure 5:**
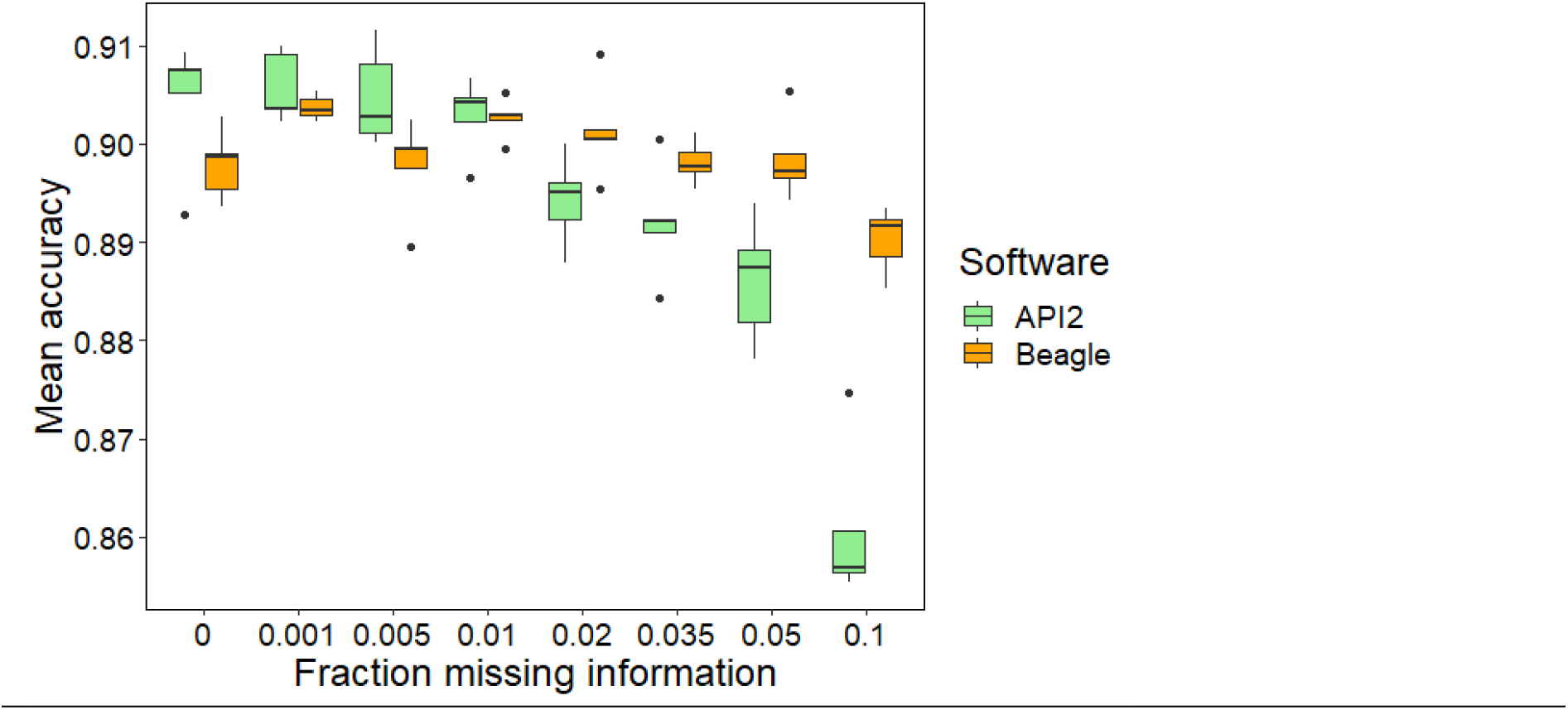
Effect of the fraction of sporadically missing genotype calls in low-density and high-density lines.

With randomly placed incorrect marker information, the accuracies for both software decreased more than for comparable fractions of missingness (compare Figure 5 and Figure 6). At 0% genotyping mistakes, accuracy for each software was 0.9 and with 10% genotyping mistakes, the accuracies were 0.53 (Beagle) and 0.58 (AlphaPlantImpute2).

**Figure 6:**
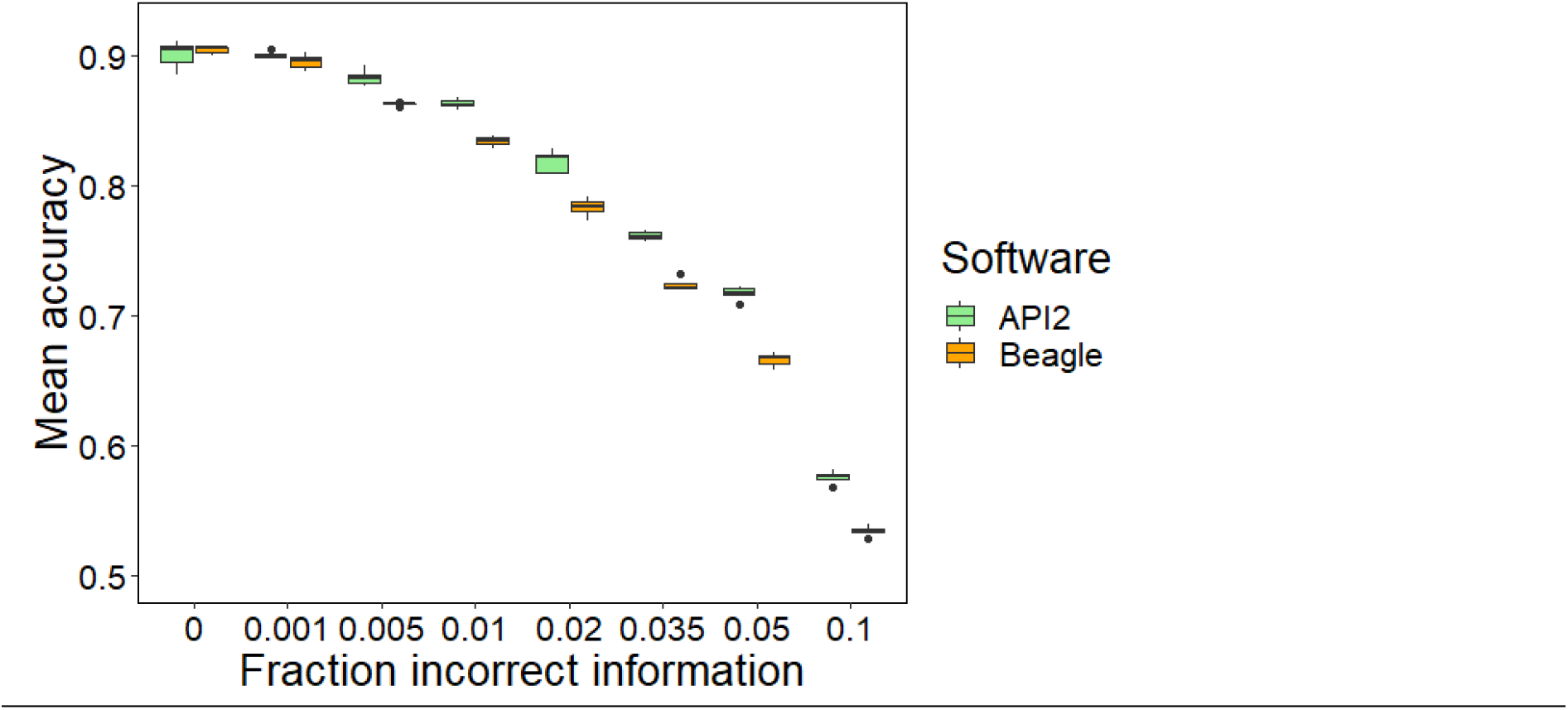
Effect of the fraction of random genotyping errors on imputation accuracy.

### 3.4 Population type

Figure 7 shows the accuracies for different parent and offspring type combinations. For bi-parental populations, imputation was done either familywise (Beagle), or the parents were specified for each line with a founders file (AlphaPlantImpute2). For open-pollinating populations based on 30 founders like OP10n5 (10 generations of open pollination at population size 5), considering the pedigree was not possible, i.e., the procedures used were those marked as ‘initial’ in Figure 1 together with optimized parameters. For multi-parent populations, 180 offspring were always imputed regardless of the population size simulated during open-pollination generations.

**Figure 7:**
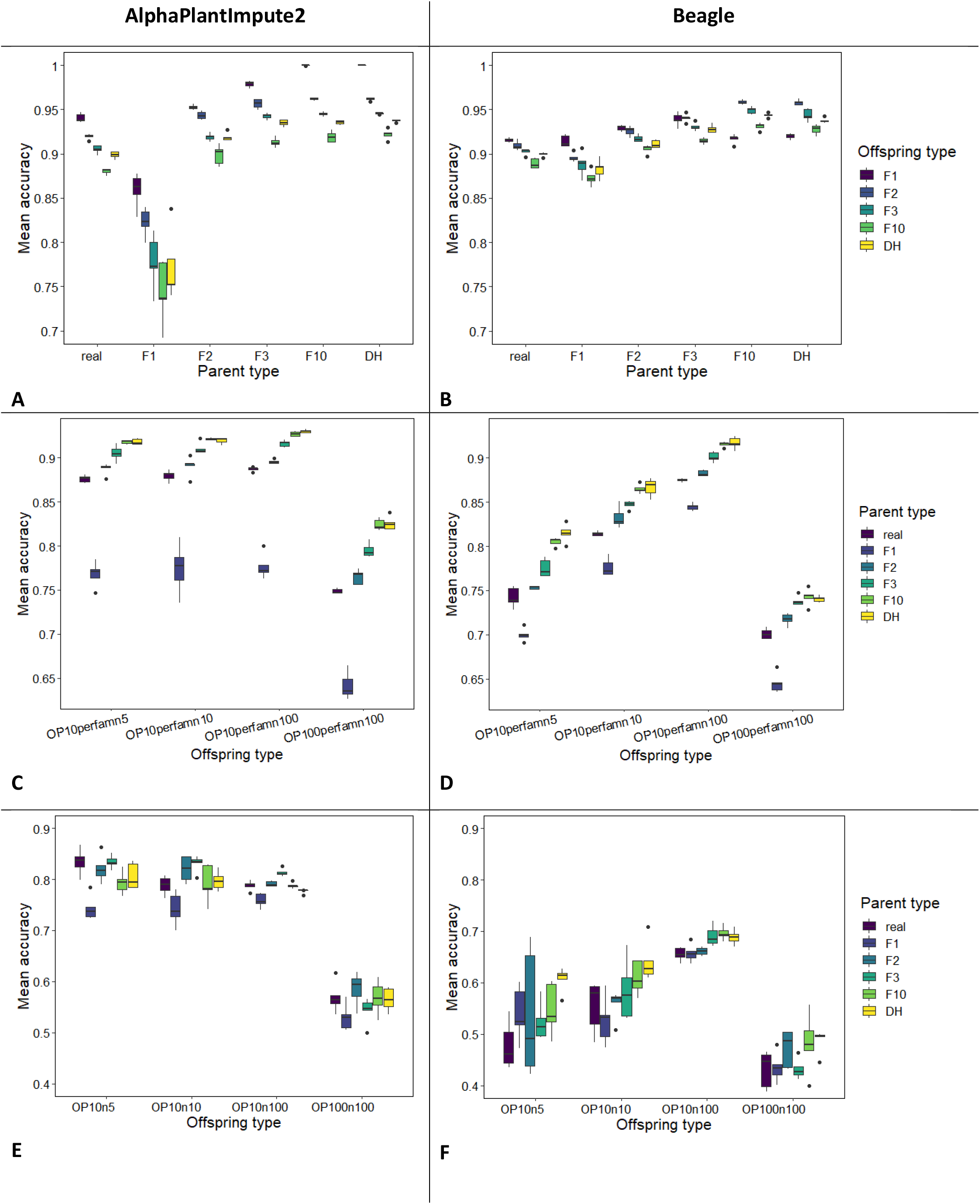
Effect of different population types of parents and offspring on mean accuracy. (A), (C) and (E) refer to imputation results with API2; (B), (D) and (F) refer to imputation with Beagle.

The imputation accuracies for Beagle and API2 increased with higher levels of homozygosity in parents (Figure 7). For offspring, the opposite trend was observed. DH offspring resulted in higher accuracies than F10 offspring. When using Beagle, the accuracies for F1 progeny was the highest for the parent types real, F1, F2 and F3 but were the lowest for the highly homozygous types F10 and DH (see plot B in Figure 7).

For open-pollinating and bi-parental intercross populations, the accuracies were better the fewer generations separated the focal population from the parents (compare OP10perfamn100 with OP100perfamn100 in (C) and (D) and OP10n100 with OP100n100 in (E) and (F) in Figure 7). A positive relationship between historic population size and imputation accuracy was observed for Beagle but not for AlphaPlantImpute2.

Generally, accuracies of AlphaPlantImpute2 exceeded or were comparable to the ones obtained with Beagle with the exception of populations based on F1 parents. API2 was able to perfectly impute F1 offspring derived from DH parents whereas Beagle achieved only a mean accuracy of 0.92 (compare plots A and B, Figure 7). Populations with DH offspring based on F1 parents are one of the traditional population types that yielded one of the lowest accuracies both with Beagle and API2 (0.88 and 0.77, respectively) (see plots A and B in Figure 7).

For further investigation, the procedures were changed so that the low-density lines were not included in the population specified to Beagle for phasing. Low-density lines were not used for the creation of the haplotype library by API2 according to the standard procedure. For investigation, we changed the API2 procedure so that low-density lines were utilized as well. Note that even though low-density lines were used for the creation of a haplotype library (API2) or phased reference set (Beagle), they were removed from those reference sets for the second step of imputation in the procedures of both software.

Excluding low-density lines from the phasing step of Beagle increased accuracies to about 0.91 for populations of DH offspring derived from F1 parents. Including low-density lines in the haplotype library creation step of API2 increased the accuracy to 0.85 (see Supplementary Figure S1new). The accuracy of API2 improved, however, at the expense of run time which increased from 76 minutes to 475 minutes.

### 3.5 Reconstruction of parents

Figure 8 shows the ability of AlphaPlantImpute2 and Beagle to impute low-density parents. Accuracies achieved with API2 are better than those achieved with Beagle for all factor level combinations. The relative advantage of AlphaPlantImpute2 was highest for the 1k low-density panel and 1 offspring per family genotyped at high-density with an accuracy of 0.89 versus 0.83 for Beagle.

**Figure 8:**
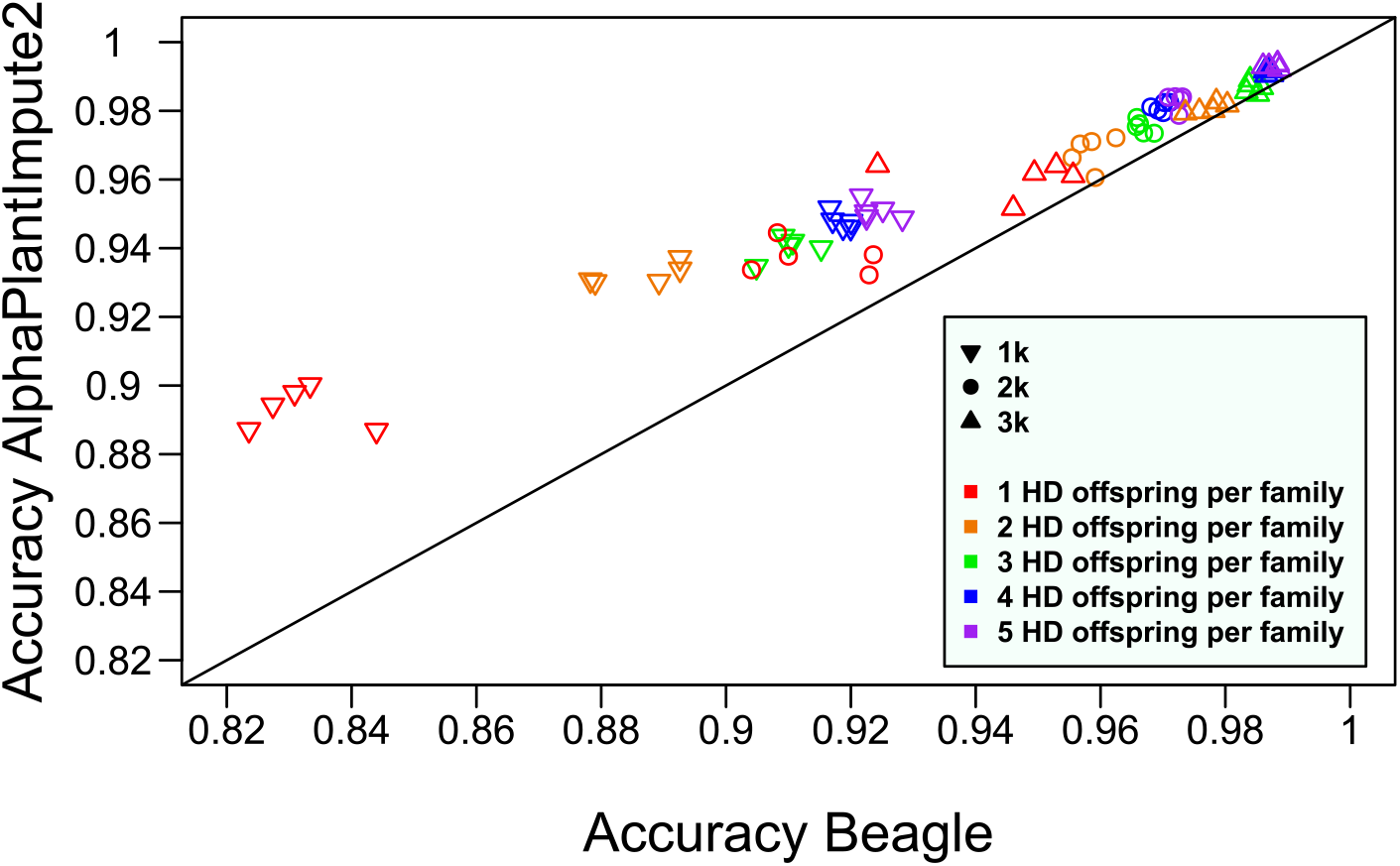
Effect of number of high-density genotyped offspring (1 to 5) and low-density panel (1k, 2k, 3k) on imputation accuracy when imputing low-density parents. Each family has five offspring lines of which some are genotyped at high-density. Accuracy is based on imputed parents only. Imputation was not done familywise.

The run time of AlphaPlantImpute2 increased by about 80 minutes for every additional line per family genotyped at high-density starting at 100 minutes. When imputing with Beagle, a decrease in run time could be observed the more lines are genotyped at high-density with 270 minutes for 1 high-density line per family and 180 minutes for all offspring genotyped at high-density. No effect of the marker number of the low-density panel on run time was found for AlphaPlantImpute2 and Beagle.

### 3.6 Family size

For all tested family sizes, the accuracies achieved when imputing with AlphaPlantImpute2 remained constant at 0.9. For Beagle, on the other hand, the imputation quality decreased the more offspring are included in a family (Figure 9). For Beagle, imputing data sets with five offspring per family (210 lines in total) took about 300 minutes whereas the run time increased to about 1740 minutes when families consisted of 50 offspring (1830 genotypes in total). For AlphaPlantImpute2, the run time increased by only about 5 minutes from 75 minutes to 80 minutes.

**Figure 9:**
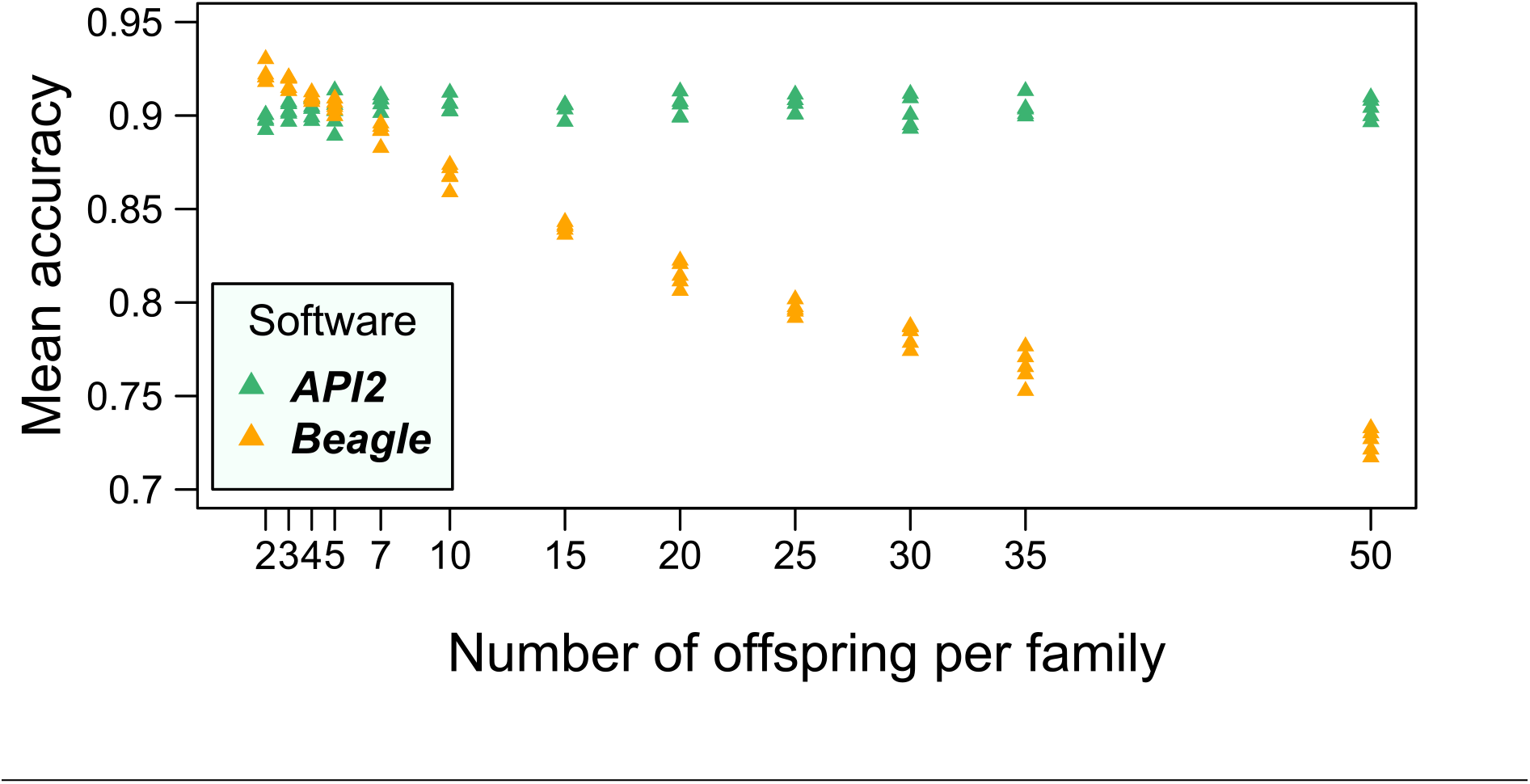
Effect of number of offspring per family on accuracy.

The procedure presented here for Beagle imputation restricts the reference set to the two parents of a family. Another difference compared to non-familywise imputation (procedure ‘2 steps’ in plot A, Figure 1) is that the genotype set to be imputed in the second step contains fewer lines since it is restricted to a family. To test whether the restriction to the two parents or the size reduction of the genotype set is responsible for the higher accuracy in our familywise imputation method compared to non-familywise imputation, we tested familywise imputation with all 30 parents in the reference. Thus, this procedure has the same size of the genotype set as our standard familywise procedure (i.e., 5) but the reference is identical to non-familywise imputation. The accuracies of this altered procedure were almost as high as with familywise imputation (Supplementary Figure S2). This suggest that the reduction of the reference set to the ancestors only has a minor effect in the familywise imputation procedure.

### 3.7 Pedigree-free tests and new methods

The Beagle procedure used in this section is the one labeled ‘2 steps’ in Figure 1 with the modification that low-density offspring genotypes are excluded from the creation of the reference set. The latter has been found to result in equal or better accuracies than if low-density genotypes were used for the reference set creation (Supplementary Figure S1). The AlphaPlantImpute2 procedure used is the one labeled ‘parents for hap lib’ in Figure 1. Thus, Beagle and AlphaPlantImpute2 considered exactly the same information at every step. The optimized parameter settings were used.

We also test two new pipelines, single-genotype imputation with Beagle 5.1 (‘single genotype imputation’), i.e., only one line is imputed at a time, and Beagle 5.1 combined with AlphaPlantImpute2 (‘Beagle + API2’), i.e., Beagle is used for phasing and API2 for imputation. For these pipelines, the ‘iterations’ parameter of Beagle was changed to 12 (default). All other parameters are identical to the optimized ones presented in Table 2 and Table 3.

Table 4 shows the accuracies achieved for all tested methods. The best accuracies for all scenarios were obtained with the Beagle + API2 method. Single-genotype imputation outperformed the 2-step procedure of Beagle. Not only was the Beagle+API2 method best in terms of accuracy, but it was also the fastest (Supplementary Table S1). To impute 5 F3 offspring per family (36 families) based on real parents, Beagle+API2 needed about 14 minutes followed by AlphaPlantImpute2, Beagle and single-genotype imputation with Beagle which took 87, 187 and 204 minutes, respectively.

**Table 4:** Mean genotype accuracies for different imputation methods testes in different scenarios. Standard error of the mean shown in parentheses. Best accuracies per scenario are underscored and in bold. LD = low-density; HD = high-density.

| parents | offspring | # offspring per family | API2 | Beagle | Single genotype imputation Beagle | Beagle (phasing) + API2 (imputation) |
| --- | --- | --- | --- | --- | --- | --- |
| real (HD) | F3 (LD) | 5 | 0.88 (0.0013) | 0.82 (0.0031) | 0.87 (0.0018) | **<u>0.89</u>** (0.0006) |
| real (HD) | F3 (LD) | 50 | 0.88 (0.001) | 0.67 (0.0059) | 0.87 (0.0006) | **<u>0.89</u>** (0.0008) |
| F1 (HD) | DH (LD) | 5 | 0.77 (0.137) | 0.85 (0.0035) | **<u>0.88</u>** (0.0032) | **<u>0.88</u>** (0.0018) |
| DH (HD) | F1 (LD) | 5 | 0.91 (0.0021) | 0.79 (0.0041) | 0.9 (0.005) | **<u>0.98</u>** (0.0011) |
| real (LD) | F3 (HD) | 1 | **<u>0.9</u>** (0.0032) | 0.87 (0.0038) | 0.86 (0.0044) | **<u>0.9</u>** (0.0043) |

AlphaPlantImpute2 performed worse than Beagle + API2 when imputing F1 lines based on DH parents (0.91 vs 0.98). Manual inspection revealed that in this scenario, API2 sets one chromosome per pair to missing in the haplotype library for the first three chromosome pairs. The other chromosome is represented correctly and chromosomes 4 to 9 are represented in the haplotype library without any missing states.

### 3.8 Beagle versions

We tried to optimize parameters for version 5.2 and 5.3 and searched for better parameters in the same ranges as done for version 5.1. With default their respective default parameters, version 5.1, 5.2 and 5.3 achieved on average accuracies of 0.23, 0.41 and 0.22 in 4.5, 3.5 and 2.2 minutes, respectively. During optimization, version 5.2 and 5.3 were found to be less responsive to parameter tuning. The effects of changing each parameter while keeping others at default are shown in Supplementary Table S2 for version 5.2 and in Supplementary Table S3 for version 5.3. Most notably, changing ‘ne’ in version 5.2 had no effect. While ‘ne’ did affect the accuracies for version 5.3, optimizing parameters for this version was generally more difficult than for version 5.1. When run with the optimized parameters found for version 5.1 as presented in Table 3, 5.1, 5.2 and 5.3 achieved on average accuracies of 0.82, 0.40 and 0.72 in 209, 190 and 213 minutes, respectively.

## 4 Discussion

### 4.1 Tuning parameters

Tuning parameters helped to improve accuracies, especially for Beagle which was developed for human data that typically show very different linkage disequilibrium and homozygosity patterns than plants. Previous work by Pook et al. (2020) on maize DH-lines showed that tuning Beagle’s parameters can yield better imputation results. The parameters declared as optimal and different from default in this study deviate even more from default than those found by Pook et al. (2020). Firstly, this might be caused by different population structures of the considered datasets. Secondly, the imputation procedure used in this study for parameter optimization differed from the one used by Pook et al. (2020). Thirdly, a much larger range for parameters than in Pook et al. (2020) was considered. This is particularly relevant for those parameters controlling the number of iterations performed, as corner solutions (i.e., better or equal than default values were at either border of the tested range) were reached in Pook et al., while we still observed improvement for a far higher number of iterations. Lastly, the number of markers and the number of lines used in this study was far lower suggesting that the importance of optimization increases the less information is available. Note that the parameters described as ‘optimized’ in this study are specific to the sugar beet data we used. Ideal parameter values for other data might be different from our findings.

Based on the observations presented in this study, we recommend using Beagle on plant breeding data only with adjusted parameters. For optimization of parameters in other data sets, we recommend to first change the value for the ‘window’ parameter so that the whole chromosome is covered if computing time allows. In a second step, the ‘ne’ parameter (effective population size) should be adjusted as this parameter has a large effect on accuracy. Plot A in Figure 2 suggests that using a precisely calculated value of the effective population size does not matter as accuracies did not change for our data set for values in the range of 1 to 250. We suggest using a value of 100 for breeding populations when a precise estimate is not available. This value is supported by a number of publications which estimate effective population sizes for crop and livestock populations much closer to 100 than to 1 million which is the default setting (Cowling, 2007; Gorssen et al., 2020; Leroy et al., 2013; Makanjuola et al., 2020; Saura et al., 2021; Zhao et al., 2021). After the publication of Pook et al. (2020), some studies have already adopted lower values for ‘ne’ (Thorn et al., 2021; Whalen & Hickey, 2020) and a few are also increasing the window size (Arouisse et al., 2020; Geibel et al., 2021; Lamb et al., 2021).

The optimized value of 4 for ‘ne’ presented in this study was chosen after the optimization process during which multiple parameters were altered at a time. We do not expect that a different value, e.g., 50, would have resulted in much different accuracies. Our proposed optimized choice is therefore somewhat arbitrary.

The parameter ‘iterations’ should be increased last since it significantly affects the run time. No negative effect of increasing the number of iterations was observed in this study. With respect to the linear increase in run time when increasing ‘iterations’ (Figure 3), a potential approach to find a good value for ‘iterations’ may be to impute the data once with default settings (i.e., 12) and a second time with any other value. This allows to approximate the additional run time for one additional iteration which can then be used to calculate the maximum number of iterations that allow imputation within the time limit. Additional iterations do not guarantee better accuracies as found by Boison et al. (2015) for earlier versions of Beagle on cattle data.

Changing parameters for AlphaPlantImpute2 did not have as drastic effects on accuracy as for Beagle which does not surprise considering that AlphaPlantImpute2 what developed for plant breeding data. This is in line with reports of Thorn et al. (2021). After extensive search, we found parameters with which AlphaPlantImpute2 performs slightly better. However, our findings do not object using default parameters.

### 4.2 Reflection on methods

It was found that Beagle’s imputation quality depends on the number of low-density genotypes provided in the imputation set (set with lines to impute) (see Figure 1, Supplementary Figure S2 and Table 4). This is because the genotype set provided for imputation impacts the haplotype cluster and library in Beagle that initializes the Hidden Markov Model and thus the effective weighting of different founder haplotypes for the imputation pipeline. In other words, imputation is not only affected by the reference set but also by the set that is to be imputed. Although the lines in the imputation set may be closely related as it is the case in our familywise approach, sibling lines do not provide additional information if the parents are included in the reference as additional crossovers compared to the parents should only introduce noise. This can be avoided with the here presented single-genotype imputation approach.

Separating an imputation problem into two steps for phasing of high-density genotypes followed by imputation of low-density genotypes was found to be advantageous for Beagle. Imputation with Beagle in two steps has been done in other studies before (Boison et al., 2015; Thorn et al., 2021). Many tools such as the here used AlphaPlantImpute2 have this separation implemented by default.

We initially used low-density lines in the phasing step of the 2-step Beagle procedure as we believed that these lines may provide some additional useful information. However, we found that excluding them is advantageous (Supplementary Figure S1).

AlphaPlantImpute2 was found to be better in most scenarios in terms of accuracy. AlphaPlantImpute2 achieved this although it utilizes less information than Beagle as it does neither accept a genetic nor a physical map. However, its advantage over Beagle diminishes the more markers are used for genotyping, the more lines are genotyped and the less homozygous the genotypes are that have to be phased. This is in line with findings by Thorn et al. (2021). Its run time is considerably higher compared to default parameters in Beagle (Figure 1). The run time for the imputation step of AlphaPlantImpute2 scales quadratically with the number of haplotypes in the reference (Thorn et al., 2021). In this study, the run time of AlphaPlantImpute2 is mainly affected by the phasing step as our standard procedure of imputation considers pedigree information thus reducing the reference to only the two parent lines.

Pedigree information improved imputation of AlphaPlantImpute2 and Beagle slightly. Pedigree information also reduced the run time for both software (see plot D in Figure 1 and Supplementary Figure S2). Note that Beagle 5.1 is not able to use pedigree information on its own but manual work to adjust the reference panels is needed. Accounting for ancestor information is a new approach that is suggested in this manuscript. For Beagle, closer investigation showed that the restriction to the two respective parents only has a minor effect on accuracy while the main improvement of familywise imputation comes from the reduced number of genotypes in the imputation set (see Supplementary Figure S2) as discussed above. This suggests that Beagle is quite good in finding the right parents from the reference which has also been shown by Pook et al. (2020). We suspect that the effect of restricting the reference set may be higher when suboptimal parameter values are used. Our findings suggest that Beagle is superior in phasing as shown by populations based on F1 parents in Figure 7. AlphaPlantImpute2 is superior in imputation as shown by populations with F1 offspring in Figure 7. Coupling the strengths of both software in our Beagle + API2 pipeline resulted in the highest accuracies and fastest run time as shown in Table 4 and Supplementary Table S1, respectively. The properties of both software may be exploited for future development of imputation software development for populations with long-range linkage disequilibrium. Note that although we presented the Beagle + API2 approach without accounting for pedigree information, utilization of pedigree information can be integrated. It should be noted that although Beagle was not the best tool in most of our tests, it is a general solution that can be applied to more different data sets than AlphaPlantImpute2 which was developed only for plant breeding.

The run times presented in this study should be interpreted as the relative advantage of one method over another. The elapsed wall clock time could be decreased further by running imputation in parallel per chromosome and per family or genotype. Single-genotype imputation with Beagle would profit the most from parallelization. Much shorter run times per CPU can be achieved by using deterministic imputation software such as FImpute (Sargolzaei et al., 2014) or AlphaPlantImpute (Gonen et al., 2018).

### 4.3 Application

Low-density genotyping of offspring may reduce genotyping costs. Together with resource saving, genotyping with a reduced marker set followed by imputation should be evaluated with regards to the effects imputed genomes have on the downstream analysis like genomic prediction or GWA studies. Moghaddar et al. (2015) report no negative effect on genomic prediction for imputation accuracies >0.95. Imputation of lines previously genotyped with a low-density array can also make those lines usable in downstream applications and further increase the size of the training population.

However, imputing low-density offspring is different from imputing progenitors. Offspring might show unique linkage at recombination points never observed before or in their sibling lines. Thus, imputing the regions around crossovers is particularly difficult. It can be mitigated by larger low-density chips. Imputing low-density progenitors has other implications. Imputation is relatively easier because the linkage pattern found in their genomes is largely passed on to the next generation genotyped at high-density. Forces like selection and drift may cause the loss of alleles or haplotypes in the current population that were present in its progenitors’ genomes (Cowling, 2013). Such lost variation is impossible to accurately impute. Supplementary Figure S3 visualizes the two problems.

For commercial breeding programs that routinely use new and denser marker chips, this raises the question for the optimal strategy to do imputation over breeding cycles. A possible solution may be to impute chronologically backwards, i.e., first the parents of the most recent generation are imputed. Then, the grandparents are imputed with the parents (grandparents’ offspring) as reference and so forth. This ensures that the haplotype segments in the reference and imputation set are as similar as possible. Boison et al. (2015) propose a similar sequential imputation strategy in which a subset of individuals with the highest genomic relationship to a fixed number of high-density individuals is imputed first and then added to the reference set to impute other individuals. Moghaddar et al. (2015) improved the imputation accuracy by only including the 20 most related animals in the reference set for every low-density animal in a study on Merino sheep. These two techniques may be extended to be applied on a chromosome level for further accuracy enhancement. Toghiani et al. (2016) investigate the imputation accuracy over several cycles when using a low-density chip and the starting generation as a fixed high-density reference. They substantially mitigated the imputation accuracy decay by updating the reference set with the high-density genotypes of top animals every generation. This is in line with our findings that genotyping high-density sibling lines in addition to parents increases the accuracy which is commonly reported in literature (Gonen et al., 2018; Hickey et al., 2012; Hickey et al., 2015; Whalen et al., 2020). If limited resources for genotyping are available, we thus propose genotyping the most promising lines if preliminary genomic breeding values are available. Alternatively, a fixed number of most related lines per parent can be genotyped starting from the most influential parent until genotyping resources are exhausted. This ensures that the reference set will be most relevant for most lines.

### 4.4 Missing and incorrect genotype calls

The effect of various proportions of missing and incorrect genotype calls on imputation accuracy was investigated. AlphaPlantImpute2 performed slightly better than Beagle in the presence of missing genotype calls while Beagle outperformed AlphaPlantImpute2 when incorrect genotype calls were induced. In practice, missing calls and genotyping errors are common (Beissinger et al., 2013; Unterseer et al., 2014) but their occurrence rates are affected by the genotyping technique and population diversity making it difficult to translate the presented findings in a general suggestion. The sensitivity of the software might change the more lines and markers are genotyped. It should be noted that positions set to missing and genotyping errors were sampled at random in this study. Functional reasons like deletions, may cause missing calls which would however result in some sites being more affected than others. Different alleles in segmental duplications, which are retained in the ancient polyploid maize genome for example, may appear as heterozygous calls in homozygous lines (Schnable et al., 2009; Unterseer et al., 2014).

Beagle provides an R2 estimate to spot problematic markers. Such poorly predicted markers could then be excluded from the data set prior to downstream applications as has been suggested by Boison et al. (2015) and Pook et al. (2020).

### 4.5 Population types

The findings of other studies stating a positive effect of the homozygosity level of parents on imputation accuracy could be confirmed (Gonen et al., 2018; Hickey et al., 2015). Those parts of the genome that are homozygous are effectively phased. Thus, the phasing step is not necessary for fully inbred parents as they are phased by design (Pook et al., 2021). A negative trend between imputation accuracy and the number of meiotic events separating parents from offspring generation has been found for open-pollinating populations and selfing populations. The more generations are separating parents from offspring, the more crossovers can break down linkage between alleles as seen in the parents. As an example, double haploid and F10 lines can both be considered fully inbred. Thus, if the homozygosity level was affecting the imputation accuracy, no difference in accuracy between F10 and DH offspring should be found. As shown in Figure 7, the contrary is the case with DH lines resulting in higher accuracies than F10 lines. This can be attributed to the fact that F10 lines show twice as many effective crossovers than DH lines. As explained above and shown in Supplementary Figure S3, imputing alleles around recombination points is imprecise.

### 4.6 Beagle versions

Beagle 5.2 performed better than version 5.1 and 5.3 when using default parameters and setting parameter ‘em’ to ‘false’ in Beagle 5.3. Changing parameter ‘ne’ had an effect for version 5.1 and 5.3 but not for version 5.2. This is because Beagle 5.2 estimates a value for ‘ne’ based on the data in the window (Browning et al., 2021) only using the specified value as a starting point for estimation. In our data set, Beagle 5.2 estimated values in the high hundreds. Based on observations with Beagle 5.1 (plot A, Figure 2), the best values for ‘ne’ are however below 250. Thus, we believed that the estimation of the effective population size may not be accurate for plant breeding data. We communicated this finding with the developer Brian Browning who then released version 5.3 in which the automated estimation of the effective population size can be turned off by setting ‘em’ to ‘false’. Although accuracies of up to 0.72 were achieved with 5.3, the best imputation quality among all Beagle versions was realized with 5.1. The ranking of the versions may depend on the population structure and marker density. It should be noted that versions 5.2 and 5.3 are developed to run much faster which will be advantageous for large datasets.

## 5 Conclusions

Our findings inform breeders seeking to obtain full high-quality data even when only a small fraction of markers is genotyped. Thus, imputation can be used to reduce costs in plant breeding. Though the results presented in this paper are based on sugar beet, the findings should be transferable to other crops as the genetic architecture of datasets should be comparable. For more fair comparisons, future work in plant breeding imputation should not use imputation software with default settings that were developed for other purposes. Our tests suggest that AlphaPlantImpute2 is a powerful tool for the imputation of plant breeding data and in particular datasets with limited genetic diversity or pedigree information. Beagle 5.1 led to competitive results when adapting parameter settings and should be more versatile on data panels with more diversity than the panel considered here. The suggested pipeline herein, involving the combination of both tools, led to the best results. Default settings of Beagle were shown to be not suitable for the here considered datasets. Among the Beagle versions, Beagle 5.1 could be optimized the most resulting in higher accuracies than 5.2 and 5.3.

## Data Availability

The genotype data and genetic map are proprietary to KWS SAAT SE. Data of the first chromosome can be obtained from the authors upon reasonable request. The R scripts for the imputation procedures are available in the GitHub repository. For testing purposes, we also provide a simple dummy dataset.

## Author contribution

TN and MG designed the experiment. TN performed the experiments, analyzed the results, and wrote the manuscript. MG contributed the raw data. TN, MG, TP, and TB discussed findings and revised the manuscript. All authors read and approved the final manuscript.

## Supplementary Material

Supplementary Figure S1: Effect of including low-density offspring in phasing step.

Supplementary Figure S2: Effect of reducing reference set to parents and imputation set to offspring lines of same family.

Supplementary Figure S3: Visualization of problem when imputing low-density offspring and low-density parents.

Supplementary Table S1: Run times of imputation methods and scenarios as shown in Table 4.

Supplementary Table S2: Accuracies and run times achieved with Beagle 5.2 with the 2-step procedure when varying one parameter while others are default.

Supplementary Table S3: Accuracies and run times achieved with Beagle 5.3 with the 2-step procedure when varying one parameter while others are default.

