## Supplementary Figure S1 for "Imputation of Low-density Marker Chip Data in Plant Breeding: Evaluation of Methods Based on Sugar Beet"

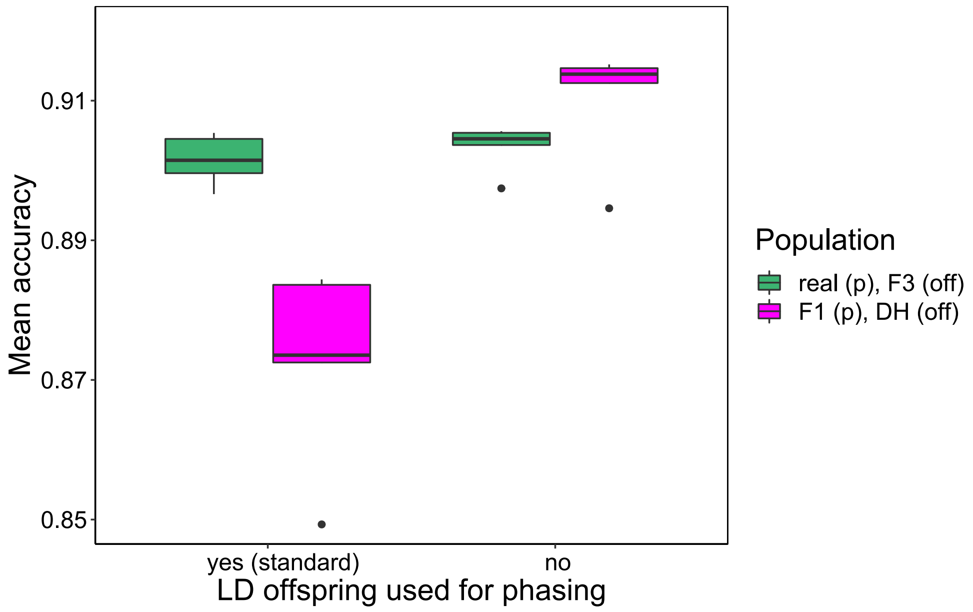


**A**

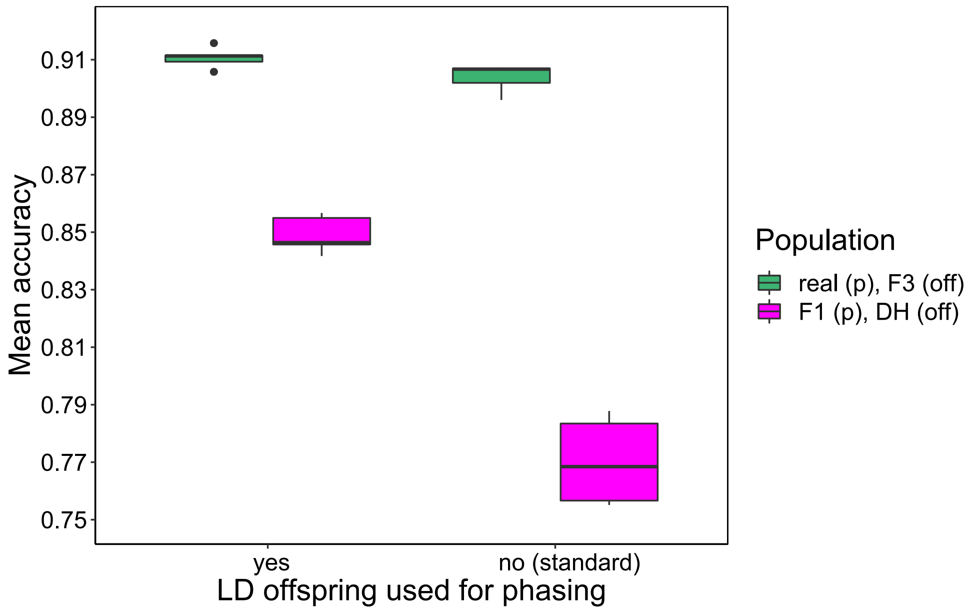


**B**

Supplementary Figure S1: Effect of including low-density offspring in phasing/haplotype library creation step of Beagle and AlphaPlantImpute2, respectively, for different population types. (A) refers to imputation with Beagle; (B) refers to imputation with API2. Note that the standard procedure of Beagle and API2 differ with regards to whether low-density individuals are used for phasing. In all cases, low-density lines are removed from the reference set/library prior to imputation.
