## Supplementary Figure S2 for "Imputation of Low-density Marker Chip Data in Plant Breeding: Evaluation of Methods Based on Sugar Beet"

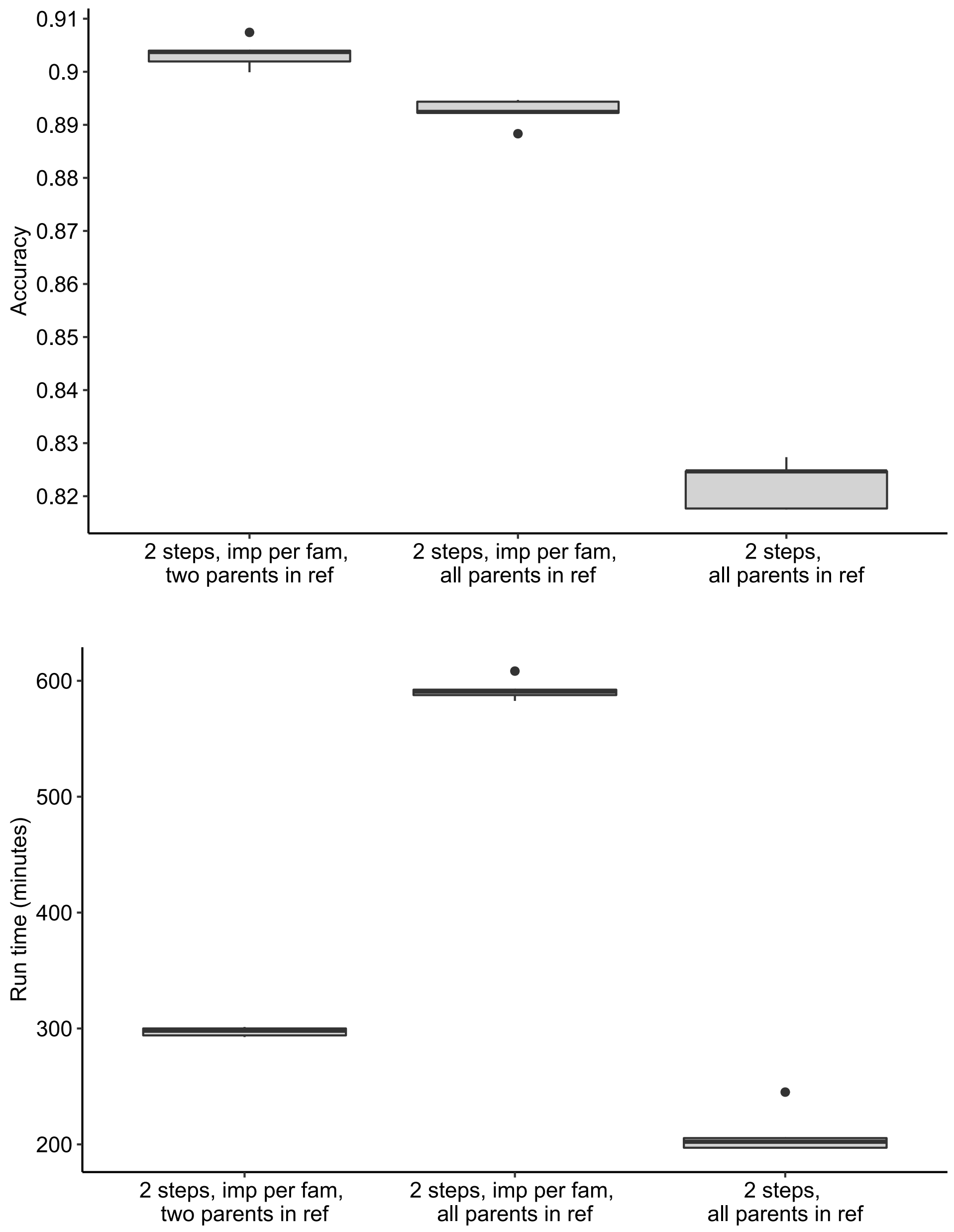


Supplementary

Figure S2: Effect of reducing number of individuals in genotype set for imputation and utilization of parent information by Beagle. The leftmost procedure is the one used as standard in this paper (familywise imputation). The second from left procedure is identical except that not only the two parents of a cross but all 30 parents are provided in the reference. The procedure on the right does not use familywise imputation (2 step procedure).
