## Supplementary Figure S3 for "Imputation of Low-density Marker Chip Data in Plant Breeding: Evaluation of Methods Based on Sugar Beet"

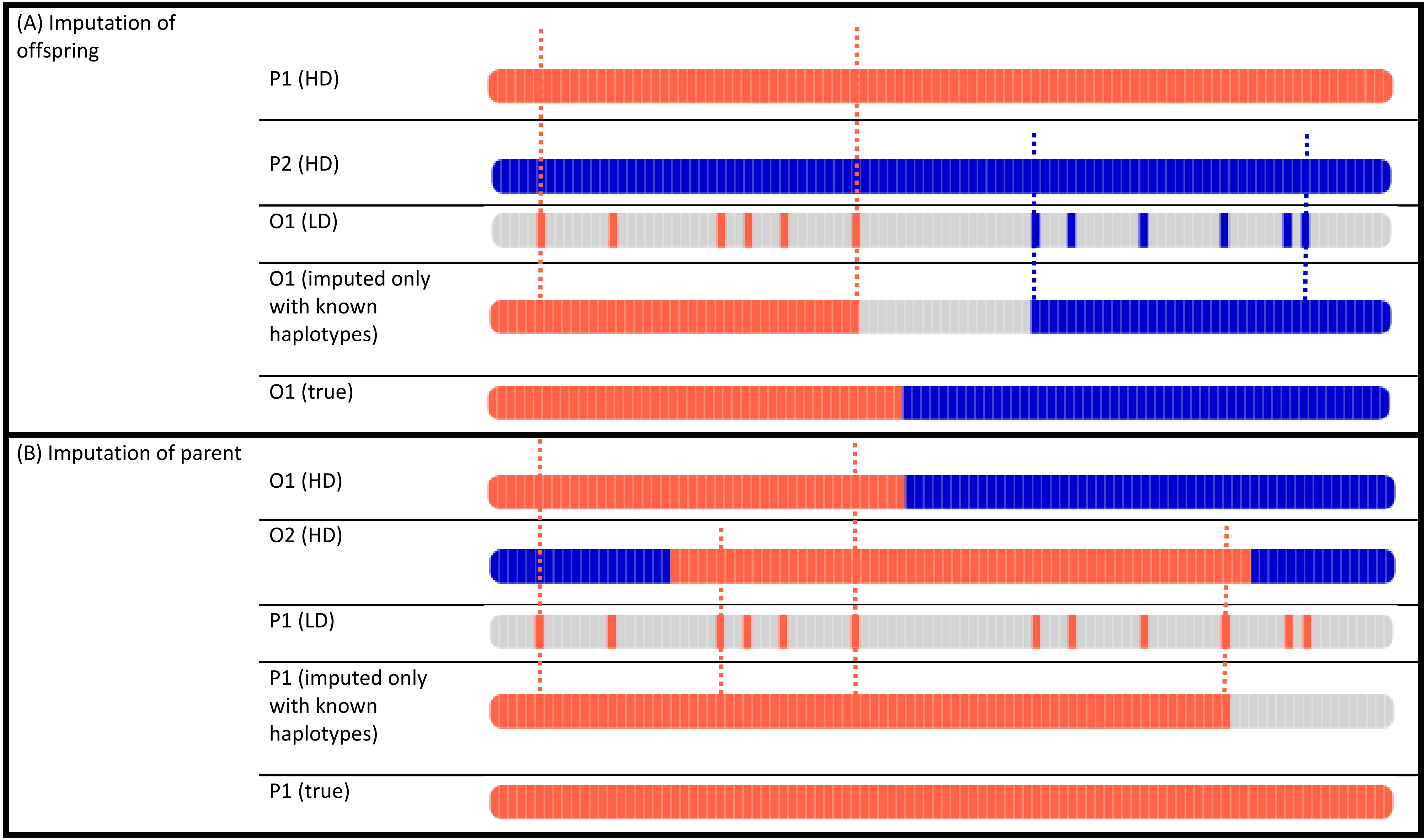
(A) Imputation of offspring
P1 (HD)
P2 (HD)
O1 (LD)
O1 (imputed only with known haplotypes)
O1 (true)
(B) Imputation of parent
O1 (HD)
O2 (HD)
P1 (LD)
P1 (imputed only with known haplotypes)
P1 (true)

Supplementary Figure S3: Offspring vs parent imputation (schematic). The species in this example is haploid. Alleles from parent 1 (P1) shown in orange and blue for parent 2. O1 and O2 are offspring of the cross P1 x P2. Imputation in this example fills the space between two alleles inherited from the same parent with the alleles from that parent. Loci surrounded by alleles from different parents cannot be imputed as the crossover can be anywhere in between.
