## Supplementary Table S1 for "Imputation of Low-density Marker Chip Data in Plant Breeding: Evaluation of Methods Based on Sugar Beet"

| parents | offspring | # offspring per family | API2 | Beagle | Single genotype imputation Beagle | Beagle (phasing), API2 (imputation) |
|---|---|---|---|---|---|---|
| real | F3 | 5 | 87<br>(0.24) | 187<br>(9.47) | 204<br>(18.38) | 14<br>(0.01) |
| real | F3 | 50 | 164<br>(0.09) | 1371<br>(68.47) | 1985<br>(95.55) | 91<br>(0.05) |
| F1 | DH | 5 | 86<br>(0.22) | 155<br>(6.38) | 260<br>(8.68) | 14<br>(0.14) |
| DH | F1 | 5 | 86<br>(0.24) | 129<br>(8.49) | 123<br>(8.73) | 13<br>(0.1) |
| real | F3 | 1 | 93<br>(0.02) | 81<br>(15.39) | 23<br>(1.38) | 7<br>(0.06) |

Supplementary Table S1: Run times of different imputation methods for selected scenarios. No pedigree information was used. The run time is reported in minutes. The standard error of the mean is reported in parentheses.
