## Supplementary Table S3 for "Imputation of Low-density Marker Chip Data in Plant Breeding: Evaluation of Methods Based on Sugar Beet"

Ranges of accuracies when varying parameters in Beagle 5.3 and using the 2-step procedure with *em=false*. The ranges of the best parameters are indicated if this was supported by a clear trend. Within the ranges, accuracy varied by about 0.02. Hyphens indicate that no trend supported stating a value as best. Variation in accuracy was thus attributed to random fluctuation.

| | default | range tested | best | accuracy (min – max) |
|---|---|---|---|---|
| ne | 1,000,000 | 1-1,000,000 | 1 - 300 | 0.22-0.44 |
| window | 40 | 3-100 | - | 0.21-0.28 |
| burnin | 3 | 1-1,000 | - | 0.21-0.23 |
| iterations | 12 | 1-20,000 | - | 0.21-0.24 |
| phase-states | 280 | 10-10,000 | 10 - 30 | 0.20-0.31 |
| imp-states | 1,600 | 10-5,000 | - | 0.21-0.22 |
| imp-segment | 6 | 0.1-700 | - | 0.20-0.22 |
| imp-step | 0.1 | 0.001-500 | - | 0.20-0.22 |
| imp-nsteps | 7 | 1-4,000 | - | 0.20-0.23 |
| cluster | 0.005 | 0.00001-50 | - | 0.20-0.23 |
| overlap | 2 | 0.01-20 | - | 0.21-0.23 |

Ranges of run time in minutes when varying parameters in Beagle 5.3 and using the 2-step procedure with *em=false*. The ranges of the best parameters are indicated if this was supported by a clear trend. Within the ranges, run time varied by about 1 minute. Hyphens indicate that no trend supported stating a value as best. Variation in run time was thus attributed to random fluctuation.

| | default | range tested | best | run time (min – max) |
|---|---|---|---|---|
| ne | 1,000,000 | 4-1,000,000 | - | 2.2-3.1 |
| window | 40 | 3-100 | - | 2.3-3.6 |
| burnin | 3 | 1-1,000 | - | 2.3-180 |
| iterations | 12 | 1-20,000 | 1 - 12 | 1.4-2908 |
| phase-states | 280 | 10-10,000 | 10 - 120 | 0.9-3.9 |
| imp-states | 1,600 | 10-5,000 | - | 2.5-4.9 |
| imp-segment | 6 | 0.1-700 | - | 2.5-5.4 |
| imp-step | 0.1 | 0.001-500 | - | 2.6-5.3 |
| imp-nsteps | 7 | 1-4,000 | - | 2.6-5.2 |
| cluster | 0.005 | 0.00001-50 | - | 2.5-3.5 |
| overlap | 2 | 0.01-20 | - | 2.5-4.0 |
